# MAPseq: improved speed, accuracy and consistency in ribosomal RNA sequence analysis

**DOI:** 10.1101/126953

**Authors:** João F Matias Rodrigues, Thomas SB Schmidt, Janko Tackmann, Christian von Mering

**Affiliations:** Institute of Molecular Life Sciences and Swiss Institute of Bioinformatics, University of Zurich, Switzerland.

## Abstract

Metagenomic sequencing has become crucial to studying microbial communities, but meaningful taxonomic analysis and inter-comparison of such data are still hampered by technical limitations, between-study design variability and inconsistencies between taxonomies used. Here we present MAPseq, a framework for reference-based rRNA metagenomic analysis that is up to 30% more accurate (F_1/2_ score) and up to one hundred times faster than existing solutions, providing in a single run multiple taxonomy classifications and hierarchical OTU mappings, for both amplicon and shotgun sequencing strategies, and for datasets of virtually any size. Availability: Source code and binaries are freely available at http://meringlab.org/software/mapseq/

## Introduction

Sequencing the DNA of microbial communities, either wholesale or after amplification of selected marker genes, has greatly advanced our understanding in many fields, including ecology, evolution and medical microbiology. Thousands of studies have generated such sequences, and more are being accrued by ever-larger projects at increasing speeds. However, the sheer amount of data and the conceptual and technical variability introduced by the wide variety of sequencing and analysis approaches pose difficult challenges for the consolidation and inter-comparability of findings, within and across studies.

The most widely used common denominator for inter-comparisons is taxonomy classification based on ribosomal RNA (rRNA), implemented in a number of software packages including RDP Classifier (Wang, et al., 2007), RTAX (Soergel, et al., 2012), MOTHUR (Schloss, et al., 2009), QIIME (Caporaso, et al., 2010), USEARCH (Edgar, 2010), VSEARCH (Rognes, et al., 2016), and NINJA-OPS (Al-Ghalith, et al., 2016). However, these packages are either restricted to previously known, validly named taxa only, are suffering from computational limitations in terms of speed and data-size, or cannot be applied in a reference-mapping mode at the scales currently needed. In addition to mapping sequences to existing taxonomies, sequences should also be mapped to *operational taxonomic units* (OTUs) – and, ideally, these OTUs should be created at various different identity cutoffs and related to each other hierarchically (Schloss and Westcott, 2011; Schmidt, et al., 2014). Such *hierarchical OTUs* (hOTUs) constitute an operational taxonomy in themselves, and enable the assessment of taxa (even uncharacterized taxa) across different studies, at adjustable levels of granularity. MAPseq enables both, by providing a fast and accurate sequence read mapping against hierarchically clustered and annotated reference sequences. In addition to the software itself, we provide a large, curated reference of full-length rRNA genes, preclustered into hOTUs at different identity thresholds, and pre-classified to taxonomic categories based on the NCBI taxonomy, and the All-species Living Tree Project dataset ((Yilmaz, et al., 2014), see Methods).

## Results

MAPseq is ten times faster than its closest competitor NINJA-OPS, and a hundred times faster than VSEARCH, on amplicon data (Fig. 1a). Despite the large speedup, the memory requirements of MAPseq are lower than those of all other tools tested here (Fig. 1b).

A routine run to analyze a typical high-throughput sequencing sample (for example a WGS metagenome of 15 million reads), requires less than three hours on a single CPU. We have also employed MAPseq in a comprehensive analysis of 646,337 metagenomic sequencing runs available in the NCBI Sequence Read Archive as of December 2016, a dataset that included both WGS and amplicon data and totaled 86TB of compressed sequence data. This computation required less than three months of runtime on a small computing cluster.

Figure 1 shows accuracy benchmarks based on placements of reads of known identity, against hOTUs clustered at 98% identity (panels d and f) as well as against taxonomic categories at genus level (panels c and e). We also tested MAPseq’s performance on reads of different lengths (panels e and f), as well as different hyper-variable regions of the rRNA gene (panels c and d). With few exceptions, MAPseq outperformed all other tools, achieving a maximum F_1/2_-score (weighted harmonic mean of precision and recall) of 0.86 in genus mapping and 0.96 in OTU mapping. The accuracy increase over existing tools is most notable in genus classification, with 30% better F_1/2_-score at 500bp long reads (Fig. 1e). The increased computational efficiency and higher accuracy are due to several innovations in the MAPseq algorithm, including improved *k-mer* counting based on a pre-clustered database, full Needleman-Wunsch alignment of high scoring segment pairs, and a sensitive algorithm to compute classification confidence; see Methods for details.

**Fig. 1.**
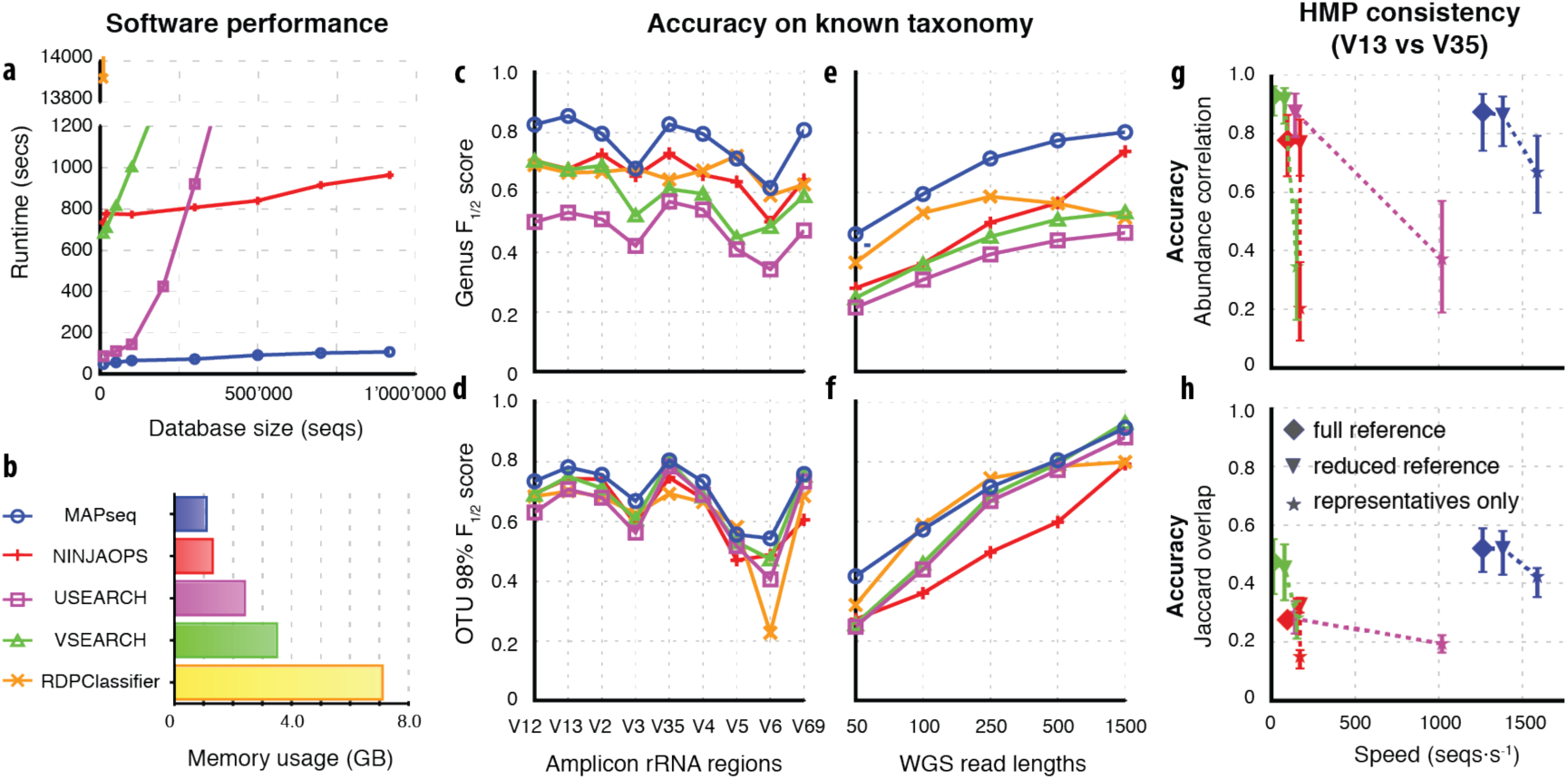
Benchmarking results on rRNA classification tasks. (a) Runtime complexity with increasing database size. (b) Maximum memory usage during benchmarking. (c, d, e, f) F_1/2_-scores for OTU (98%) or taxonomy (genus) classification, testing fragments of varying lengths or from different rRNA regions. (g, h) Concordance tests on 2,194 Human Microbiome Project (HMP) samples, sequenced twice for both the V1V3 and V3V5 regions; (g) Pearson correlation of OTU abundances, and (h) overlaps in terms of OTUs identified.

For all methods, the F_1/2_-score tends to improve with increasing read lengths (Fig. 1e, f). The F_1/2_-scores of reads originating from hyper-variable regions within the rRNA gene were significantly better (up to 20%) than those of reads originating from random positions in the rRNA gene with similar read lengths (mimicking non-targeted study setups). This latter observation is as expected (due to the higher information content of variable regions), and it is particularly pronounced for reads of smaller lengths covering a single hyper-variable region.

Strikingly, we find that even small reductions in the comprehensiveness of the reference dataset, by removal of near-identical sequences, can affect the accuracy for all methods tested (Fig. 1g, h; Fig. S2). The F_1/2_-score is lowered by as much as 0.1 when mapping to a representative reference, as compared to a full reference. This shows that making reference datasets less redundant for runtime reasons has a significant trade-off cost in terms of mapping accuracy.

For an independent test of classification accuracy, we took advantage of a large data collection for which the very same samples had been subjected to two independent sequencing runs, using different regions of the rRNA gene. Here, any good analysis framework should report strongly correlated results. This was based on a subset of the Human Microbiome Project (HMP) (Huttenhower, et al., 2012) consisting of 2,194 samples where sequencing runs for both V1-V3 and V3-V5 regions were available. As shown in Fig. 1g, the correlations of abundances of mapped OTUs between sequencing runs are found to be fairly high for most methods, however MAPseq achieved by far the best trade-off in terms of speed versus accuracy. We observed a similar trend in terms of the fraction of shared OTU identifications (Jaccard overlap, Fig. 1h): MAPseq resulted in the highest overlap (median=0.52), followed by VSEARCH (median=0.47), NINJA-OPS (median=0.27), and USEARCH (median=0.28). Finally, we investigated the effect of using a different method for the *ab initio* clustering of reference sequences into OTUs. Using reference OTUs obtained with UCLUST, a widely adopted method, resulted in lower overall abundance correlations (median=0.62) and Jaccard overlaps (median=0.33) (Fig. S3). This reduction in accuracy is independent of the software subsequently used for the mapping (Fig. S3).

In summary, MAPseq outperforms state-of-the-art methods dramatically in terms of speed, while also providing a more accurate and consistent approach, both for targeted as well as for untargeted metagenomic sequences. MAPseq can be applied routinely and quickly to individual samples, but it can also be used to analyze very large and diverse sequence collections in a consistent way, because of its use of a common reference. MAPseq is open-source software implemented in C++, and uses multiple cores when available. Both the software and the corresponding reference database are available for download at: http://www.meringlab.org/software/mapseq/.

## Methods

### Reference dataset

A dataset of publicly available full-length small subunit ribosomal RNA sequences was compiled from the databases RefSeq and NCBI Genbank. Sequences were aligned using INFERNAL v1.0.2 with three rRNA models for Bacteria, Archaea and Eukarya, respectively, as available in the ssu-align package (Nawrocki, et al., 2009). Sequences with a negative structure score, or an alignment score less than 40, were discarded. After alignment, insertions were removed and sequences were trimmed between two well-conserved alignment columns (Bacteria: 107 and 1408, Archaea: 142 and 899, and Eukarya: 629 and 1547), such that all sequences had the same aligned length. Sequences not covering the full length between the trimmed positions were filtered out, as were sequences with more than 5% unknown (“N”) nucleotides. Chimeric sequences were removed by running UCHIME against a reference set of sequences confirmed by at least three independent studies and falling within the same OTUs (defined at 98%). The final reference set consisted of 918’803 sequences. The reference set used for the actual MAPseq mapping process is composed of the unaligned sequences, including the insertions that had been removed during alignment.

**Figure S1.**
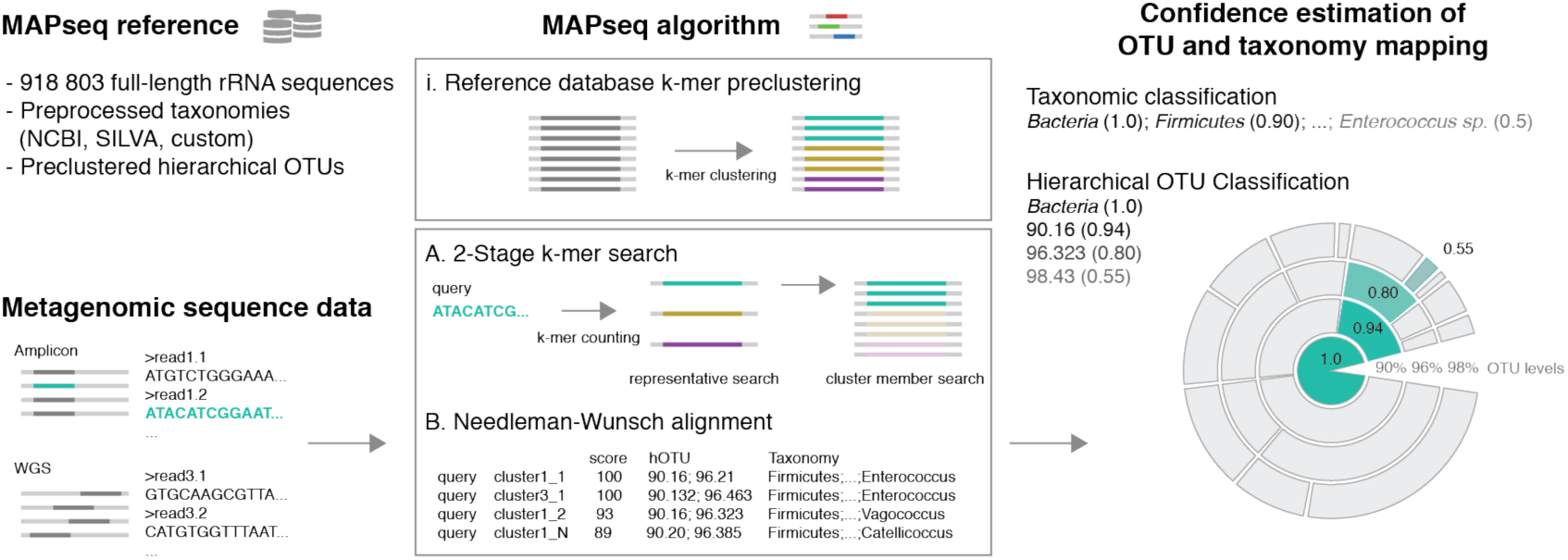
MAPseq algorithm and reference. Together with the MAPseq software, a large full-length ribosomal RNA reference dataset is provided along with pre-computed hierarchical OTUs and taxonomic assignments from NCBI and SILVA. The algorithm requires a k-mer pre-clustering of the reference dataset which is provided for the MAPseq reference and can be built for custom, user-provided references. The classification of a query sequence begins with a 2-stage k-mer search; first, the representatives of k-mer clusters are searched, and then the members of the top clusters are searched. Finally, the sequences are fully aligned and confidences are estimated for each taxonomic level.

### Reference taxonomy

NCBI taxonomies for the reference dataset were assembled either by referring to the annotated taxonomy for sequences extracted from the Refseq database, or by scanning for GenBank entries annotated to be culture collection strains. This taxonomy reference set comprised 89,315 sequences, covering 15,810 species, 3,637 genera, 1,493 families, 800 orders, 404 classes, and 185 phyla. Another, independent taxonomy annotation was obtained from SILVA Living Tree Project (LTP) (Yilmaz, et al., 2014) by mapping the sequences annotated in the SILVA LTP database to the MAPseq reference set. The set of sequences with a trusted taxonomy was defined as the *gold* set and was used to predict the taxonomy of the remaining, un-annotated sequences.

### Hierarchical OTUs

The set of aligned sequences in the Archaea, Bacteria and Eukarya reference datasets were separately clustered using HPC-CLUST (Matias Rodrigues and von Mering, 2014) down to 90% sequence identity. Each sequence was then assigned to an OTU at five different identity cutoffs: 99%, 98%, 96% and 90%, yielding 144’596, 92’862, 51’904, and 19’957 OTUs, respectively.

### MAPseq algorithm

MAPseq achieves a big advance in efficiency and accuracy at searching databases of highly similar sequences, by improving upon the k-mer counting approach used in many other sequence search tools such as BLAST or USEARCH. To achieve this efficiency, the reference dataset is pre-clustered into clusters of sequences. This pre-clustering is made available to MAPseq together with the actual sequence data and the taxonomic information. At runtime, MAPseq uses a 2-stage k-mer search – first to the k-mer cluster representatives, and second to the cluster members – to effectively reduce the number of sequences to be searched exponentially and to significantly improve the memory requirements of the program.

### K-mer pre-clustering

Sequences in the datasets to be used with MAPseq are pre-clustered on the basis of the number of shared k-mers, in two phases. The first sequence in the dataset is used as the seed for the first cluster, afterwards, every sequence in the dataset is iteratively compared on the basis of shared k-mers to the existing cluster representatives and are either assigned to one of the existing clusters or used as the representative for a new cluster. Sequences are added to existing clusters when the sequence being considered shares at least 80% of its k-mers with a cluster representative. The first phase of pre-clustering is complete once all sequences are assigned to a cluster or used as representatives for new clusters. In the second phase, all cluster representatives are kept from the first phase, but the membership assignment for the other sequences is recalculated, this ensures that sequences assigned to a cluster at some stage can be assigned to another cluster representative to which they are more similar. This situation can occur because cluster representatives can be created at any point after sequences have already been earlier assigned to other clusters in the first phase. Pre-clustering of our dataset consisting of 918’803 sequences results in 56’169 k-mer clusters which implies a 16 times reduction in number of sequences that need to be searched in the first stage of the k-mer search step.

### Mapping of sequence reads to the reference dataset

When a query is first searched against the database, the number of shared k-mers between the query and one representative for each cluster is computed. In a second step, the number of shared k-mers is computed between the query and all of the members belonging to the clusters that ranked highest in the previous step. Finally, the top hits are aligned to the query sequence. Dynamic programming is used to identify the set of identical segments yielding the largest alignment score, followed by alignment of the regions between these segments using the banded Needleman-Wunsch algorithm. The ends of the alignment are determined using the banded Smith-Waterman algorithm with drop-off such that the alignment is not extended if the alignment score drops to more than 20 below best score found to that point. The best match is taken as the best guess for taxonomy or OTU assignment, and the remaining hits are used to estimate the confidence in the assignment at each level.

### Estimating assignment confidence

MAPseq achieves higher accuracy by assigning a mapping confidence to better control false classifications. There are two different types of false classifications: i) misclassifications due to missing sequences in the reference dataset, and ii) incorrect assignments among the existing sequences in the database. These two types of errors are controlled for independently. The first case is controlled by computing a confidence on the basis of identity cutoffs previously optimized for each level for a given taxonomy and reference dataset. The identity confidence is computed using the following formula:

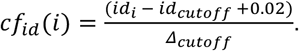

Where *id_i_* is the identity of the query compared to the reference sequence *i, id_cutoff_* is the identity cutoff, and *Δ_cutoff_* is a parameter that controls the strength of the effect of identity differences on the confidence; these two parameters were optimized previously for each taxonomic level. The additional 0.02 was added so that the *id_cutoff_* term matches the OTU cutoffs in OTU mapping.

The second case is controlled using the formula described below that weighs the score of the top hit against subsequent hits to different taxa:

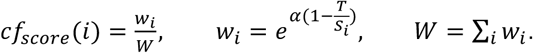

Where *S_i_* is the alignment score of reference sequence *i* in the top hit list, *T* is the score of the best aligned sequence, *w_i_* is the weight of sequence *i, W* is the sum over all weights, *α* is a parameter controlling the influence of lower scoring hits. In essence, the higher the difference in scores between the top hit and the closest second-best hits to any conflicting taxonomy/OTU, the better the confidence placed in the assignment. This automatically solves the problem of sequence reads originating in highly conserved parts of the 16S rRNA sequence, because top hits will tend to have similar scores even when belonging to conflicting taxonomies. The final confidence is calculated as the minimum between the two confidences:

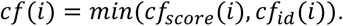

Another advantage of this approach is that higher confidences are automatically computed for lower taxonomic levels, since second-best hits will necessarily have higher score differences at lower taxonomic levels.

### Validation of sequence read mapping

Benchmarking of the taxonomic classification and OTU mapping performance was done using as a starting point a set of nearly full-length 16S/18S ribosomal subunit sequences compiled from the Genbank database. The taxonomy from the All-Species Living Tree Project (LTP) (Yarza, et al., 2010) which comprises a set of manually curated sequence and taxonomies was used as the gold standard in our benchmarks. For the OTU mapping, sequences were clustered using HPC-CLUST (Matias Rodrigues and von Mering, 2014) after alignment with the INFERNAL aligner (Nawrocki, et al., 2009), or with UCLUST.

To benchmark the accuracy of MAPseq and other classification tools commonly used in metagenomic data analysis (RDP Classifier 2.6, NINJA-OPS 1.5, VSEARCH 2.4.0 and USEARCH 9.2.64) we generated a benchmark query and reference dataset from our full-length rRNA gene dataset in which the true OTU assignments or taxonomies were known. We avoided benchmarking trivial cases when the query and database sequence were nearly identical, by excluding from the benchmark reference any sequences originating from the same species or OTU (at 99% identity cutoff) as any of the query sequences. In addition, we evaluated the ability of different methods to correctly ignore sequences that had no representative in the reference database by generating the reference set such that in the best case only 50% of the query sequences actually had a representative of their genus or OTU (at 98% identity cutoff). This test set was generated by randomly selecting one sequence for each genus from the full set while respecting the previous conditions.

Confidence thresholds per method were chosen as the thresholds yielding maximum F_1/2_-scores averaged over the scores obtained for different read lengths and different rRNA hypervariable regions. For USEARCH and VSEARCH, which output no confidences, we used the difference above the identity cutoff for each level as a measure of confidence, for example when a query had 98.1% identity to a reference sequence then it had 0.1 confidence when mapping to an OTU at 98% identity cutoff.

The query sets were generated by segmenting the selected sequences into different lengths (50bp, 100bp, 250bp, 500bp, and 1500bp) and at different positions incrementally in steps of half the segment length. For example, 50bp segments were generated starting at positions (1, 25, 50, 75, …) and 100bp were generated starting at positions (1, 50, 100, …). For the hypervariable regions, the sequences were trimmed at positions matching the *Escherichia coli* 16S rRNA after alignment with INFERNAL. The hypervariable regions were selected according to (Schloss, 2010) as follows: V2 (100bp to 337bp), V3 (357bp to 514bp), V4 (578bp to 784bp), V5 (784bp to 986bp), V6 (986bp to 1045bp), V1V2 (28bp to 337bp), V1V3 (28bp to 514bp), V3V5 (357bp to 906bp) and V6V9 (986bp to 1491bp).

The same test reference dataset and queries were used with all methods tested.

### Computation speed and memory benchmarks

The computation speed and memory benchmarks were performed by running each tool with 8 threads on the same dedicated computer on: (Figure 1a) the Human Microbiome Project sequencing run (700016012) consisting of amplicon raw data targeting the V3V5 region of the 16S rRNA gene to different subsamples of the MAPseq reference database consisting of 918 803 sequences, and (Figure 1g,h) on 10% subsampled set of all raw reads found in 2,194 HMP samples for which both V1-V3 and V3-V5 sequencing runs existed.

USARCH could not be run on reference datasets larger than 650’000 sequences due to memory usage limitations of the 32bit version, the only freely available version. The following commands were used for running the benchmarks:

### USEARCH

`usearch9.2.64 -usearch_local input.fa -db mapseqref.udb9.2 -threads 8 -id 0.90 -strand both -blast6out usearch.output`

### VSEARCH

`vsearch -usearch_global input.fa -db mapseqref.fa -threads 8 -id 0.90 - strand both -blast6out vsearch.output`

### NINJA-OPS

`python NINJA-OPS/bin/ninja.py -d 0 -p 8 -z -i input.fa -b mapseqref`

### MAPseq

`mapseq -nthreads 8 input.fa mapseqref.fasta`

### RDP Classifier

java -Xmx10g -jar rdp_classifier_2.6/dist/classifier.jar classify -t mapseqref.rdpdb/rRNAClassifier.properties -o rdpclass.output input.fa

### Concordance analysis of independent V1V3 and V3V5 sequence data of the Human Microbiome Project

Raw 16S rRNA V1V3 and V3V5 amplicon sequencing data and sample metadata of the Human Microbiome Project (HMP) were downloaded from the NCBI Sequence Read Archive and the HMP data depository (hmpdacc.org). Chimeric reads were detected using UCHIME (Edgar, et al., 2011) in both *de novo* mode and with a custom in-house reference database of non-chimeric sequences; reads labeled as chimeric by both approaches were removed from further analyses. Filtered reads were then aligned to a 16S rRNA model using INFERNAL (Nawrocki, et al., 2009) and pruned to the respective alignment flanking positions for the V1V3 and V3V5 primer sets, as described above. Reads that did not align to these regions, that had too many gaps within flanking positions or that were not observed in at least two samples independently at a 5bp error tolerance were removed from further analyses. Moreover, all biological samples were removed that did not contain at least 1,000 sequences of both V1V3 and V3V5. After these filtering steps, there remained 2,194 samples containing a total of 17,890,946 (V1V3) and 26,627,383 (V3V5) sequences.

Sequences were assigned to OTUs by reference mapping. Consistency between V1V3 and V3V5 data from the same biological samples was assessed as direct per-sample Jaccard similarity (i.e., the overlap of reference OTUs at 97% called in both V1V3 and V3V5 sequencing of the same sample) and as the Pearson correlation of abundances of OTUs shared between both sequencing sets per sample. Reference mapping was performed on three different reference databases: One including all full-length sequences for each OTU (full reference, Figure 1g,h), one with 30% randomly picked representatives per OTU (reduced reference) and one including a single random representative for each OTU (representatives only). Speed estimates are based on reduced input files (random 10% of the sequences for each sample) and tool parameters as in the previous section, while consistency (Jaccard similarity, Pearson correlation) was computed on all input sequences with 40 to 80 threads on a larger workstation.

## Supplementary Results

### Accuracy reduction when using representative reference datasets

A common practice used in ribosomal RNA marker gene analysis is to make the reference database non-redundant or to even use only representatives of each OTU. In Fig. S2 we show that this practice leads to up to 0.1 less F-score than using the full reference set (Fig. 1f). This reduction in F_1/2_-score was not limited to MAPseq but was also observed for all other tools tested.

**Figure S2.**
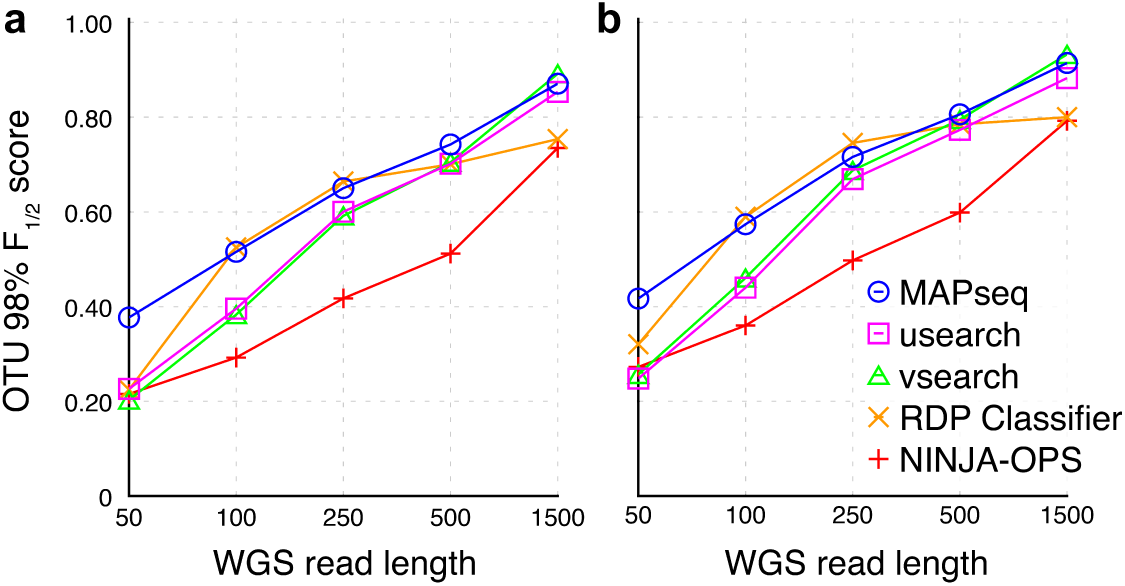
F1/2 scores for sequence reads of different lengths, mapped to OTUs at 98% cutoff using: a) non-redundant/representative reference dataset and b) the full reference, identical to Fig. 1f and included here for direct comparison.

### HPC-CLUST based OTUs outperform UCLUST OTUs in mapping consistency

We chose a subset of HMP metagenomic data in which the same samples had been sequenced twice, targeting two different regions of the rRNA gene (V1 to V3 and V3 to V5); these were in total 2,194 samples. Abundance correlations for mapped OTUs at 97% identity between sequenced subregions per sample were very high (median=0.83, mean=0.73; Figure S2a) when using MAPseq and our reference OTU clustering. When using MAPseq to map to the same reference (but pre-clustered by UCLUST, the tool used in the GreenGenes and SILVA databases) the correlation was lower (median=0.62, mean=0.59), and when using UCLUST (v1.2.22q) to map the sequences the correlations were yet significantly lower both when mapping to our reference (median=0.39, mean=0.41) as well as to the UCLUST-clustered reference (median=0.22, mean=0.29; this latter approach corresponds to the default for “closed-reference OTU picking” in the widely used QIIME pipeline). We observed a similar trend in the fraction of identified OTUs common to both pairs of sequencing runs for the same sample (Figure S3b): MAPseq mapping to the MAPseq reference resulted in the highest overlap (median=0.46, mean=0.43), followed by MAPseq mapping to the UCLUST-clustered reference (median=0.33, mean=0.31). Using UCLUST resulted in a much-reduced overlap between OTUs common to pairs of sequencing runs when mapping to OTUs (median=0.30, mean=0.29) and when mapping to UCLUST representatives (median=0.23, mean=0.22).

**Figure S3.**
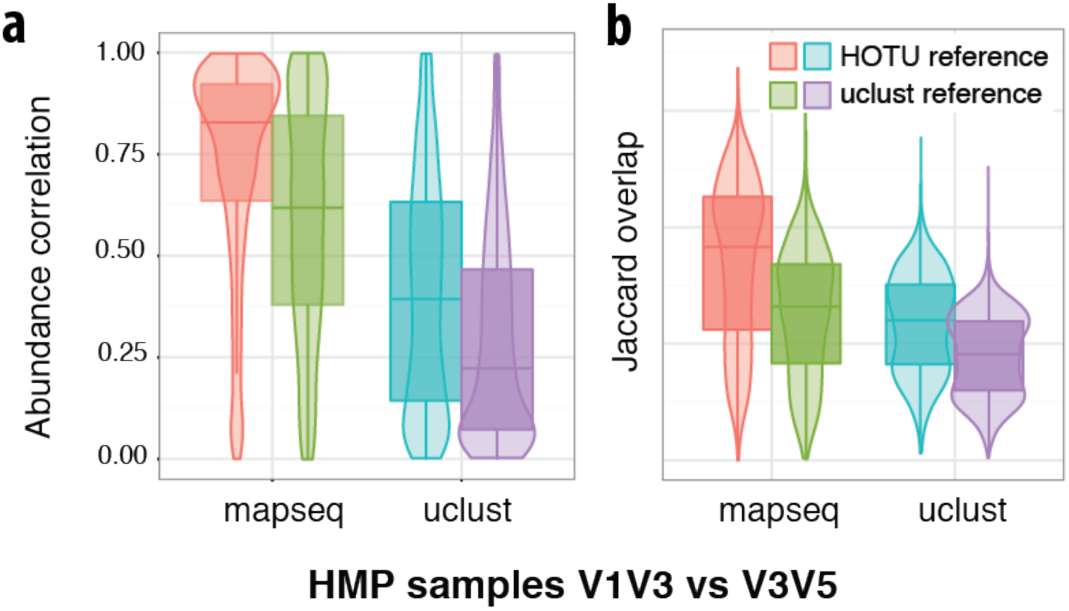
Abundance correlations and Jaccard overlaps for OTUs identified in pairs of sequencing runs of identical HMP samples targeting two different ribosomal RNA regions (V1-V3 and V3-V5). The same reference dataset was used with two different clustering approaches: hierarchical clustering using average-linkage computed with HPC-CLUST, and a UCLUST clustering.

Finally, we compared these two OTU clustering methods to *de novo* clustering of the metagenomic samples. OTU set compositional consistency between reference-mapped partitions and their respective *de novo* counterparts was checked by calculating the Adjusted Mutual Information (AMI, (Vinh, et al., 2009)) between the respective OTU sets (e.g., comparing MAPseq against an average linkage pre-clustered reference with an average linkage *de novo* clustered partition). Ideally, the results of *de novo* clustering and reference-mapping strategies should correspond to each other – if consistent sequence processing and clustering algorithms are applied. However, this correspondence has recently been called into question for UCLUST (Westcott and Schloss, 2015). To test this, we quantified partition similarity in terms of OTU composition (per-sequence cluster membership) as *Adjusted Mutual Information* (AMI); AMI values of 1 indicate perfect partition identity, and AMI values of zero indicate random compositional agreement as expected by chance. For the HMP dataset, we observed that MAPseq against an average linkage (AL) OTU reference provided a good correspondence with AL *de novo* clustering (AMI=0.83). This indicates that MAPseq reference mapping, although computationally much more efficient, may indeed approximate hierarchical AL *de novo* clustering, which is arguably still a gold standard in marker gene processing. In contrast, UCLUST mapping against a UCLUST-clustered non-redundant reference (the default method in QIIME, see above) showed much lower agreement with *de novo* UCLUST clustering (AMI=0.66). UCLUST performed better against our AL OTU reference (AMI=0.75), but was outperformed by MAPseq against a UCLUST reference (AMI=0.79). Thus, both reference pre-clustering and choice of mapping tool have a strong effect on consistency, and UCLUST was clearly outperformed on both, by MAPseq mapping and by AL clustering.

